# Decomposing the subclonal structure of tumors with two-way mixture models on copy number aberrations

**DOI:** 10.1101/278887

**Authors:** An-Shun Tai, Chien-Hua Peng, Shih-Chi Peng, Wen-Ping Hsieh

**Affiliations:** Institute of Statistics, National Tsing Hua University, Hsinchu, Taiwan, R.O.C.

## Abstract

Multistage tumorigenesis is a dynamic process characterized by the accumulation of mutations. Thus, a tumor mass is composed of genetically divergent cell subclones. With the advancement of next-generation sequencing (NGS), mathematical models have been recently developed to decompose tumor subclonal architecture from a collective genome sequencing data. Most of the methods focused on single-nucleotide variants (SNVs). However, somatic copy number aberrations (CNAs) also play critical roles in carcinogenesis. Therefore, further modeling subclonal CNAs composition would hold the promise to improve the analysis of tumor heterogeneity and cancer evolution. To address this issue, we developed a two-way mixture Poisson model, named CloneDeMix for the deconvolution of read-depth information. It can infer the subclonal copy number, mutational cellular prevalence (MCP), subclone composition, and the order in which mutations occurred in the evolutionary hierarchy. The performance of CloneDeMix was systematically assessed in simulations. As a result, the accuracy of CNA inference was nearly 93% and the MCP was also accurately restored. Furthermore, we also demonstrated its applicability using head and neck cancer samples from TCGA. Our results inform about the extent of subclonal CNA diversity, and a group of candidate genes that probably initiate lymph node metastasis during tumor evolution was also discovered. Most importantly, these driver genes are located at 11q13.3 which is highly susceptible to copy number change in head and neck cancer genomes. This study successfully estimates subclonal CNAs and exhibit the evolutionary relationships of mutation events. By doing so, we can track tumor heterogeneity and identify crucial mutations during evolution process. Hence, it facilitates not only understanding the cancer development but finding potential therapeutic targets. Briefly, this framework has implications for improved modeling of tumor evolution and the importance of inclusion of subclonal CNAs.

## Introduction

Cancer, a dynamic disease, is characterized by unusual cells with somatic mutations. These mutations are caused by environmental factors accumulated during an individual’s lifetime; this accumulation of mutational events results in a large degree of genetic heterogeneity among cancer cells. The intratumor heterogeneity causes difficulties in devising personalized treatment strategies.

To decipher intratumor heterogeneity, understanding how cancer evolves is a key step. The hypothesis for the somatic evolution of cancer was proposed in the 1970s [1]. It states that all tumor cells descend from a single founder cell, and cells with some advantageous mutations become more competitive than normal cells for growth and clonal expansion. This hypothesis could also be formed through random drift. Gradually, subsequent clonal expansion occurs, and the tumor evolves into an organization of multiple cell subpopulations. Understanding clonal evolution in cancer is one of the goals of cancer medicine [2]. Presently, sequencing technology enables performing a large-scale molecular profiling of tumors to comprehend cancer development and determine disease progression. However, the process of evolution is not directly observed because tissues for measuring somatic mutations are typically obtained from patients at a single time point. Thus, the ancestral relationship among tumor subclones have to be inferred, and this is closely related to a well-studied problem, phylogenetic tree reconstruction. To construct a phylogenetic tree, the mutations in each cancer cell should be measured to infer evolutionary relationships among various cells. For addressing this concern, the current technology of single-cell sequencing seems appropriate [3, 4]. However, this technology is not widely used because of some technical limitations and financial considerations [5]. Most studies on tumor evolution rely on DNA sequencing technology with a bulk tumor containing genetically different cells. Therefore, the cellular prevalence of each subclone have to be measured through the relative read count information of the variants.

Single-nucleotide variants (SNVs) and copy number aberrations (CNAs) are widely used data types to study tumor evolution. Recently, studies inferring the population structure and clonal architecture have either focused on SNVs according to variant allele frequencies (VAFs) or on CNAs with read counts obtained through DNA sequencing [6, 7]. Methods for either type of data can adopt the other type of data to improve their reconstruction, and most methods have developed corresponding computational tools.

The first category of method reconstructs models with only SNV data. AncesTree and clonality inference in tumors using phylogeny (CITUP) are the representatives of this category, and they build models based on heterozygous SNV to study cancer progression, assuming that the copy number is two [8, 9]. To relax the assumption of the normal copy number status, many studies have included CNAs to correct the baseline [10-13]. For instance, Pyclone is one of the clonal inference approaches, and it applies a hierarchical Bayes binomial distribution to model allelic counts [13]. This approach applies a Dirichlet process prior on group mutations and infers the posterior distribution to estimate the cellular prevalence, which is the fraction of cancer cells harboring a mutation.

Unfortunately, the aforementioned algorithms only considered abnormal copy number states but do not infer the clonal structure of copy number changes. If we do not account for clonal evolutionary architecture, the estimation of CNAs would be inaccurate and just reported as an average of the CNAs of all tumor subclones. Hence, in contrast to the SNV-based models, some studies focus on subclonal CNA heterogeneity [7, 14-18]. They recognize that subclonal CNAs could technically improve the analysis accuracy. THetA is one of the most popular tools for subclonal copy number decomposition [7]; it searches all possible combinations of copy numbers across all segments and applies the maximum likelihood approach to infer the most likely subclonal structures. However, THetA has an identifiability concern, such that several solutions of subclone structures and copy number status levels can explain the read-depth information equally well [15, 16].

Integrating other data, such as single-nucleotide polymorphisms, to jointly analyze tumor progression is a solution to the identification problem. The methods developed on the basis of these integrated data types constitute another category of cancer subclone reconstruction approaches [14-18]. In 2014, Oesper et al. modified THetA to THetA2, which designs a probabilistic model of B-allele frequencies (BAFs) to solve the identification problem and simultaneously improves the efficiency of the algorithm [14]. Furthermore, PyLOH resolves the identifiability problem by integrating CNAs and loss of heterozygosity (LOH) within a unified probabilistic model [15]. PyLOH aims at determining the contamination from normal cells and evaluating tumor purity, which is the fraction of tumor cells within a tumor tissue. Instead of tumor purity, MixClone improves PyLOH with a more delicate measurement of tumor progression, the subclonal cellular prevalence (SCP) [16]. The major concept of PyLOH and MixClone is to use the Poisson and binomial models simultaneously to analyze the read depth and BAFs.

Most of the above mentioned methods that reconstruct the process of copy number evolution assume heterozygous SNV sites within chromosome segments. This assumption facilitates the decomposition of clonal CNAs, but it ignores segments without any somatic SNVs. Therefore, to more effectively address this concern, we developed a new algorithm, called CloneDeMix, which considers subclonal copy number changes when inferring the clonal evolutionary structures. It requires only the read-depth information of loci of any sizes no matter SNVs are included or not. The input can be a predefined segment of the chromosome or simply a single nucleotide locus. CloneDeMix is a two-way clustering model that clusters each locus into an appropriate copy number state and a most likely clonal group. The procedure can simultaneously evaluate all loci and regions. The algorithm uses information from all samples and loci simultaneously to infer clone progression and can efficiently reduce the identification bias. The flowchart of CloneDeMix is demonstrated in Fig 1.

**Fig 1.**
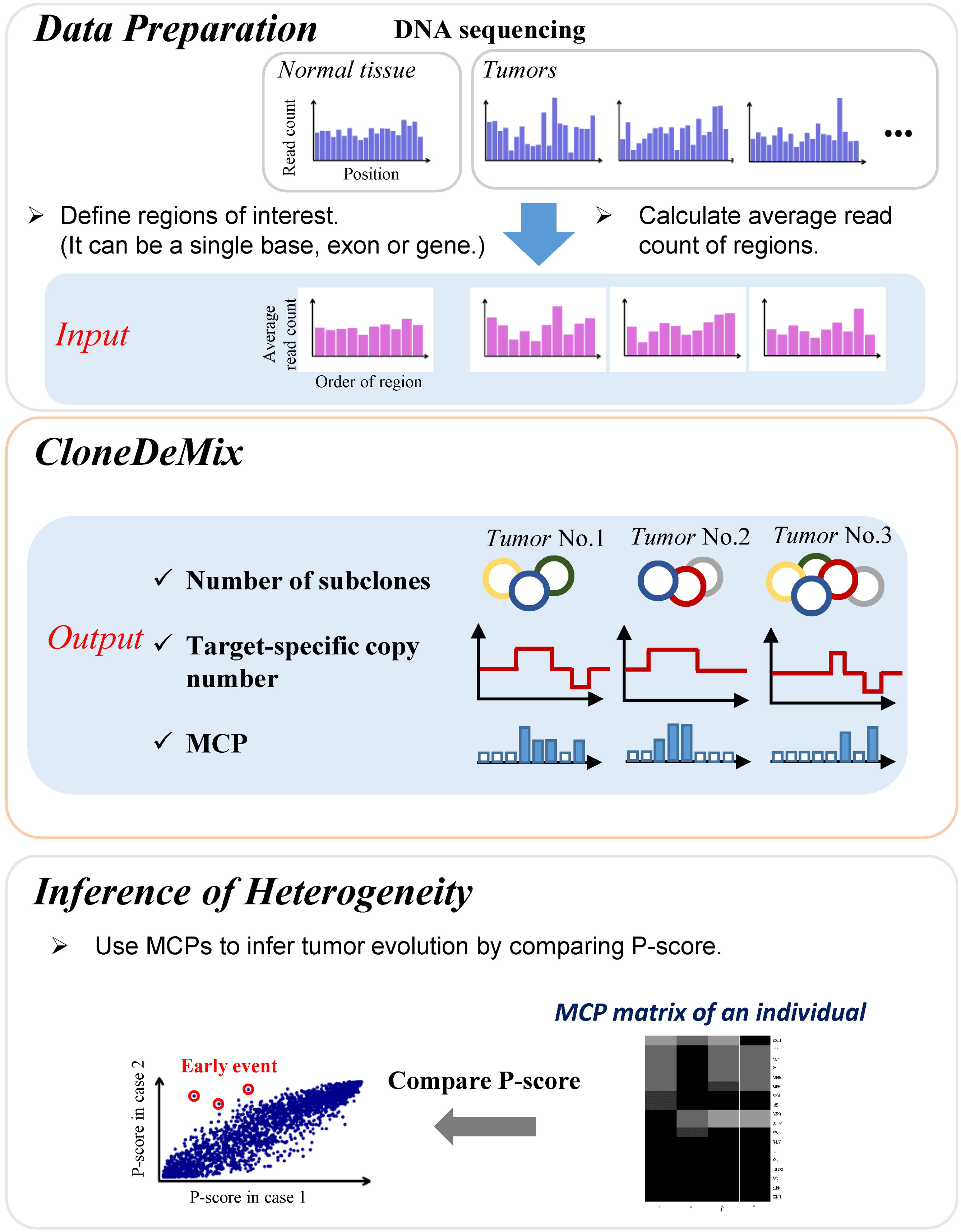
Flowchart of CloneDeMix. Our approach includes three main steps, data preparation, running CloneDeMix, and inference of tumor heterogeneity.

In this study, we demonstrated the performance of the algorithm with simulation data and applied it to a head and neck cancer dataset from The Cancer Genome Atlas (TCGA) and primary esophageal squamous cell carcinoma (ESCC) [19]. The simulation demonstrated the accuracy of clone identification and subclonal copy number change detection, particularly in early mutational events, which could be the candidate of driver mutations. The specificity of the copy number detection exceeded 98%, and the sensitivity was nearly 93.5%. These simulations support that our approach can successfully identify the copy number mutation and deconvolute its amplification or deletion state from the clonal architecture.

Our results obtained for 75 paired normal–tumor samples recapitulated most of the findings reported in head and neck cancer [20-23]. The novel subclonal CNAs have also been identified, and their subclonal structure has been shown to facilitate the discovery of driver mutations for advanced tumor progression. Furthermore, we provide evidence for the association between tumor heterogeneity and metastasis. A large heterogeneity tends to promote tumor metastasis. To sum up, CloneDeMix demonstrated ability to accurately identify subclonal CNAs and clarify intratumor heterogeneity. It is useful complement to other methods for cancer evolution studies.

## Methods

### Two-way Poisson mixture model

We delineated the structure of cellular evolution based on two concepts: SCP and mutational cellular prevalence (MCP), as shown in Fig 2. The SCP is defined as the fraction of cells that are relatively homogeneous and carry the same set of mutations. The MCP is defined as the fraction of cells that carry a certain mutation. The SCPs can be added to match the MCPs according to the evolutionary structure of subclones (Fig 3A). The evolution matrix, an upper triangular matrix, in Fig 3A provides information on the ancestral relationship among the subclones. There are five subclones in this toy example and their relationship is shown in the evolution tree in Fig 3A. The percentages indicate the corresponding SCPs. In this evolutionary structure, six mutations create five subclones. For example, locus A exists in every tumor subclone because of its presence at the top of the tree. Hence, the MCP of this locus can be calculated as the sum of all SCPs. By contrast, locus G is a later mutation and only exists in the leaf subclone C4. The corresponding MCP is equal to the SCP of C4.

**Fig 2.**
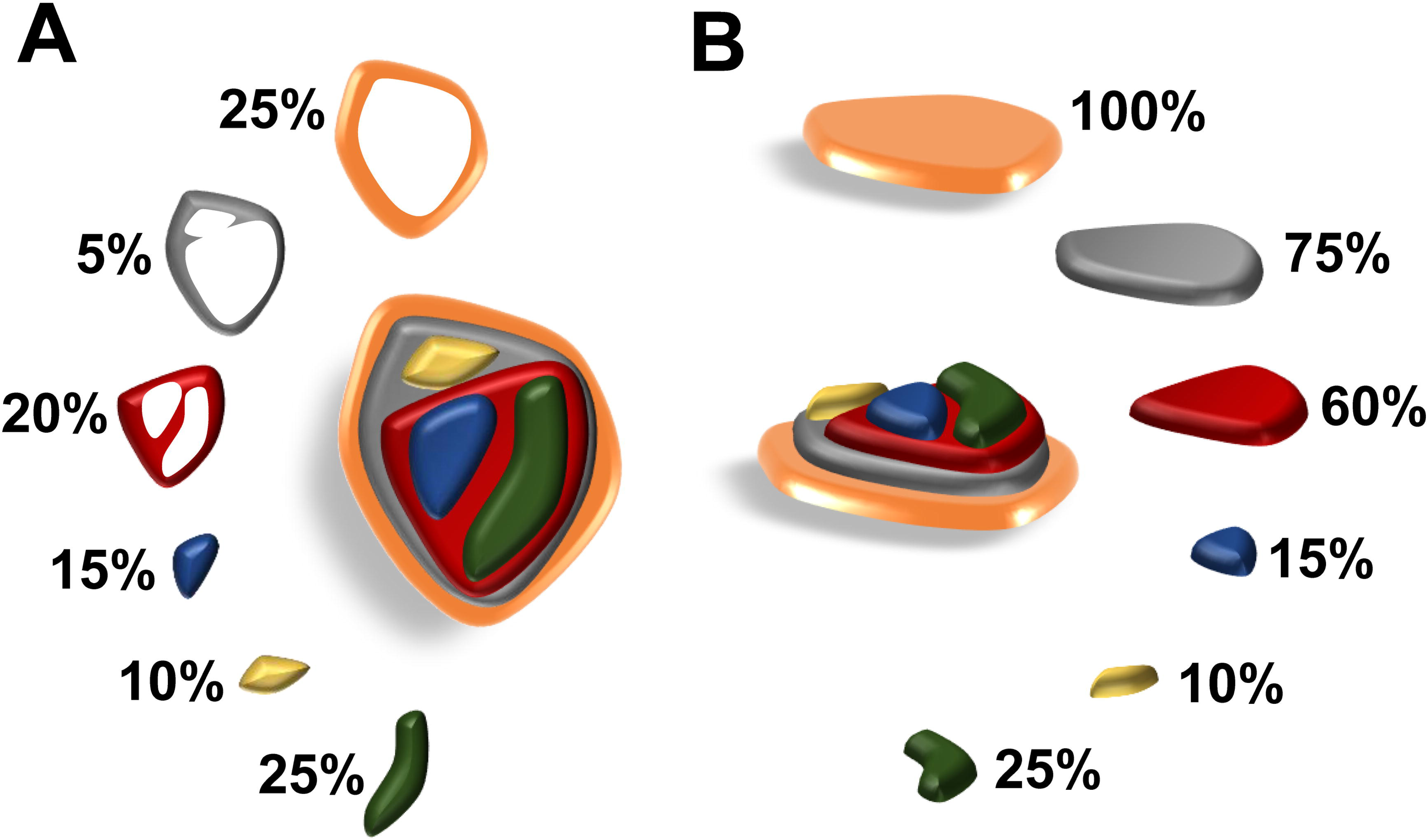
Illustration of SCP and MCP. A tissue has two decompositions. Panel (A) provides an overhead view that divides the cells into several disjoint groups according to their mutations. The cells in the same group are relatively homogeneous and carry the same set of mutations. The size of a group or the fraction of cells is called the SCP. In contrast to the SCP, panel (B) demonstrates the MCP, which is defined as the fraction of cells carrying a certain mutation.

**Fig 3.**
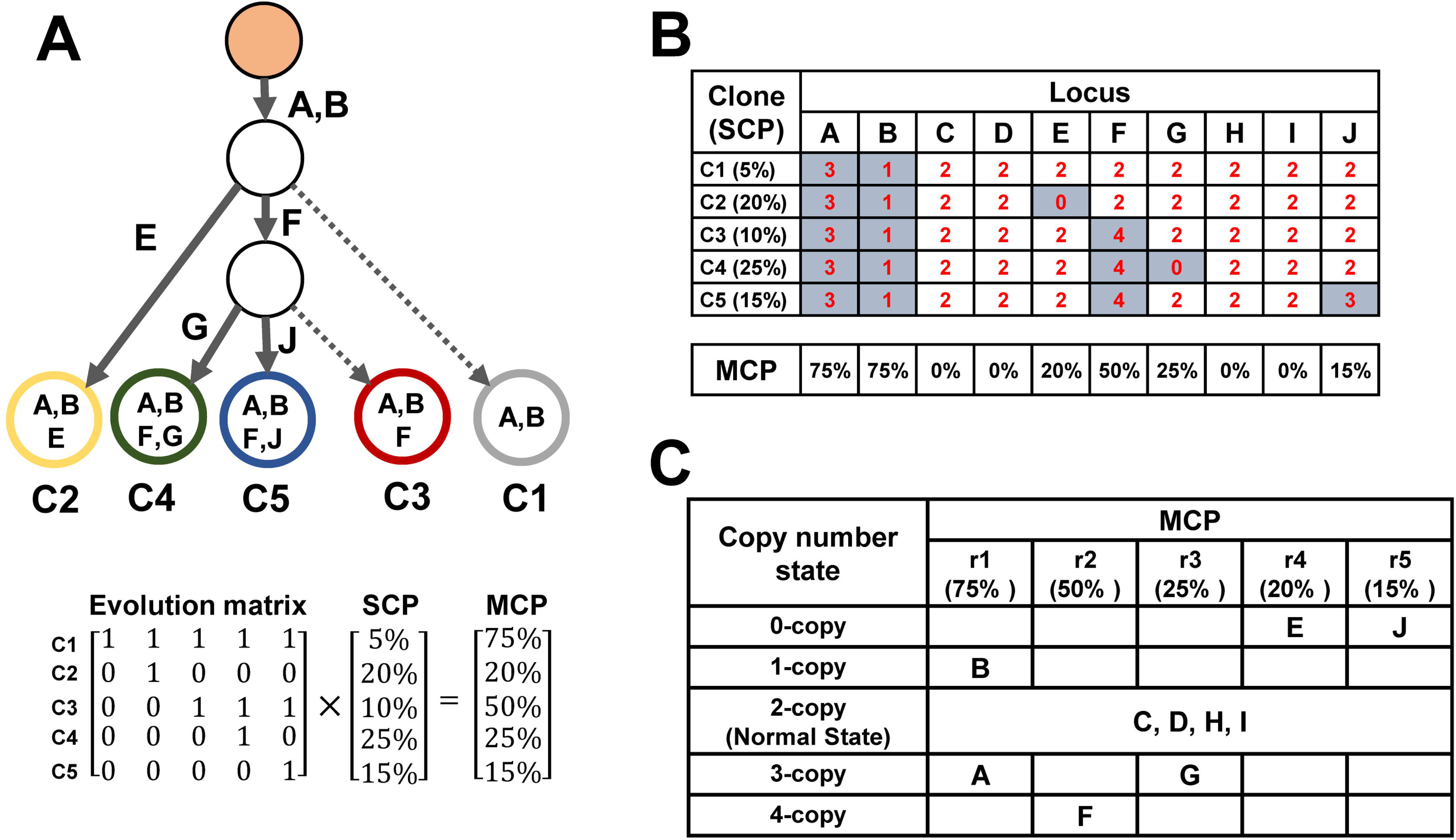
Two-way mixture model for inferring tumor progression by using copy numbers. (A) A toy example for tumor progression of five distinct subclones. Six of the ten loci (A, B, E, F, G, and J) have gained or lost copy numbers, and the remaining loci (C, D, H, and I) show no copy number change. The mutation in each locus forms a new subclone. MCPs can be determined by multiplying the SCP and evolution matrix. (B) The copy number status of each locus is listed in the table, and the MCP of each locus is listed under the table. (C) Each locus belongs to one of the 21 clusters in CloneDeMix. The columns represent five MCP levels, and the rows represent five copy number states considered in the example.

The read depth of each locus is proportional to the copy number and MCP. To delineate the read depth of each somatic copy number variant into its copy number state and MCP, this study proposes a two-way mixture model (CloneDeMix). Any locus in a sample has only two states, namely normal and mutated states; the proportion of both types differs across different loci. For example, locus E shows copy number changes in subclone C2 but not in the other subclones (Fig 3B). Hence, all other subclones comprise the normal allele for locus E. Furthermore, locus F has copy number changes in C3, C4, and C5; hence, it is classified as normal in subclones C1 and C2. CloneDeMix clusters all loci according to their copy number state and MCP. As shown in Fig 3C, all loci in this case are classified into five copy number states and simultaneously into five MCP levels. This results in 21 groups because we could not distinguish the MCP levels for the loci of two copies. The MCPs for the five MCP groups are unknown and have to be estimated. Thus, CloneDeMix provides the copy number and MCP for each locus.

The input in CloneDeMix is the read depth of each analyzed locus. When the locus represents a segment, such as an exon or a predefined amplicon, the average read depth is adopted. Let *X_i_* be the read depth of locus *i* or the average read depth rounded to the closest integer in region *i*, and assume that it follows a two-way mixture Poisson distribution.

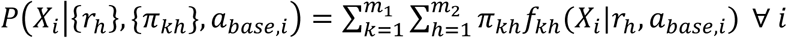

Each component *f_kh_*(*X_i_*) in the model represents the distribution of read depths sampled from the k-th and h-th groups of the copy number state and MCPs, respectively. The read count for each combined group is specified as a Poisson distribution; the mean of this distribution is proportional to a function of the mutated copy number and the MCP. It is specified as

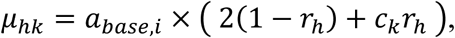
where *r_h_* is the MCP for the h-th group, *c_k_* is the copy number of the k-th copy number group, and *a_base,i_* is a normalization number for locus *i*. The corresponding mixture weight is denoted as *π_kh_*. Without further evidence, the copy number of the normal cells can be considered to be two in CloneDeMix. The number of groups for copy numbers and cellular proportions are pre-specified as m_1_ and m_2_, respectively; we select m_1_ and m_2_ according to the Akaike information criterion (AIC).

### Estimating MCPs and copy number by using expectation–maximization algorithm

The parameters of CloneDeMix include the normalization constants *a_base,i_*, MCPs **r** = {*r_h_*}, and weights **Π** = {*π_kh_*}. The plug-in estimator of *a_base,i_* is estimated from the paired normal sample of each tumor sample. Because all samples are assumed to be globally normalized, and the sample-specific variation is removed before the analysis, the read depth of locus *i* in the normal sample represents an unbiased estimator of the mean read depth in the tumor sample when the copy number is two. Hence, we use half of the read depth of locus *i* in the paired normal sample as the estimator of *a_base,i_*. In case of no paired normal sample, we suggest taking half of the sample mean across all existing normal samples to estimate *a_base,i_*. All other parameters are estimated using the expectation–maximization (EM) algorithm to approximate the maximum likelihood estimation (MLE).

We introduce a sequence of latent binary variables for locus *i*. Variables 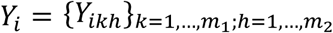 take the value of 0 or 1, indicating the memberships of the copy number and MCP groups for locus *i*. If *Y_ikh_* = 1, then 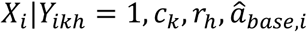 has the following distribution

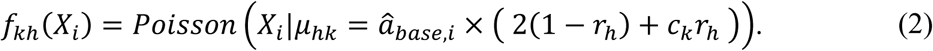

A complete form of the conditional distribution is

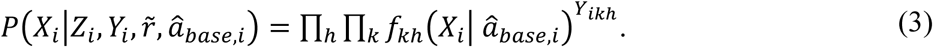

According to the mixture model construction, the probability of *Y_ikh_* = 1 is *π_kh_*. Specifically, 

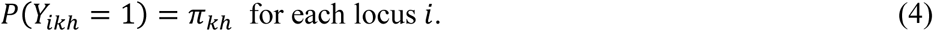

Hence, the density functions of 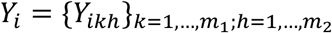 follow multinomial distributions with probability functions

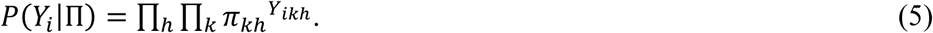

According to the definition of conditional probability, the joint density function of *X_i_* and *Y_i_* can be written as follows:

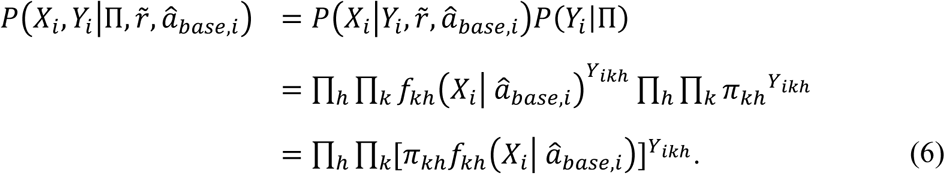

The log likelihood of Π and 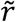 is

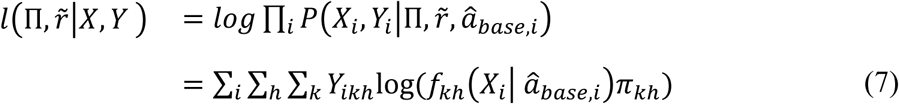

Because there is no closed form for the maximum likelihood estimator of Π and 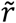, we adopted the EM algorithm to determine the MLE. The EM algorithm iteratively maximizes the expected log likelihood in two steps: E and M steps.

The E step of the EM algorithm determines the expected value of the log likelihood over the value of the latent variable Y, given the observed data X and current parameter value Π = Π^0^ and 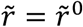. Thus, we derive the following equation:

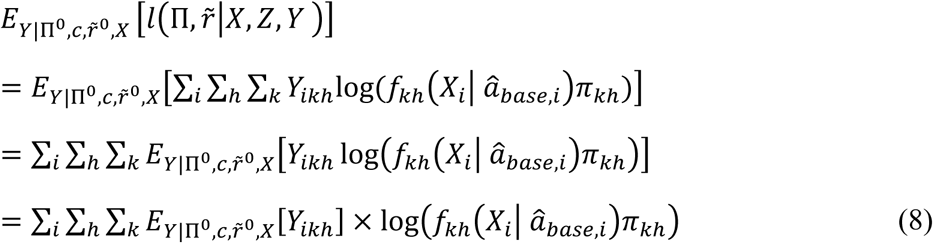

According to the definition of *Y_ikh_*, 

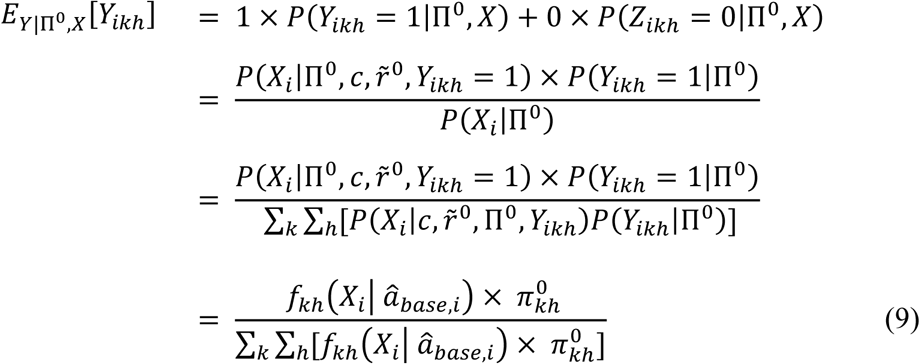

Let 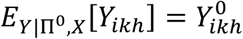 and substitute it into equation (8); with some arrangement, we obtain

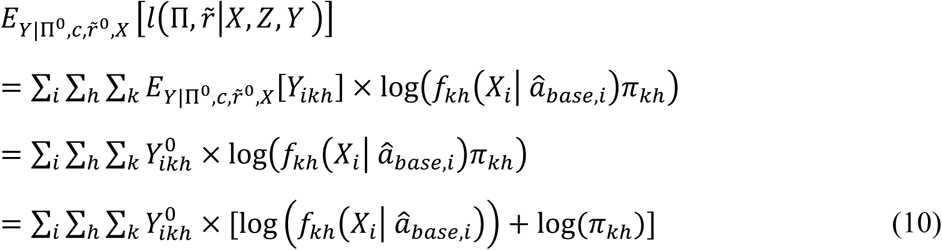

The M step of the EM algorithm maximizes equation (10) over Π, 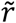 to determine the next estimates (e.g., Π^1^ and 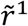). The maximization over Π involves only the second term in equation (10):

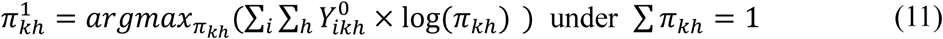

The solution is 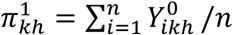. The maximization of 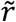 concerns the first term of equation (10), and the solution has no closed form. Numeric algorithms, such as the Newton–Raphson method, are required to solve the equation. We used the Newton–Raphson method with the R function *optim*(), and the iterative algorithm for 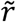 is

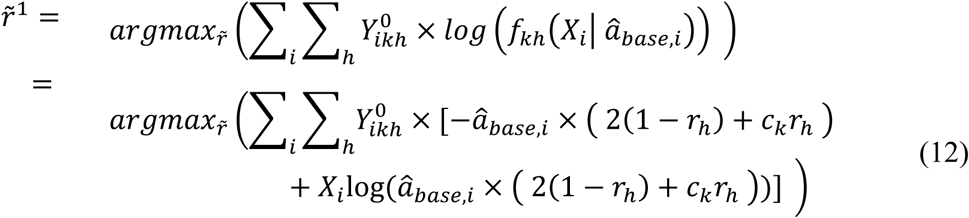

The solutions 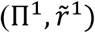 are substituted into equation (10) to replace 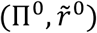. The expectation is then rewritten as 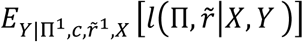. The algorithm continues iteratively to maximize the expectation of the log likelihood.

### Determining the order of copy number variants

Based on the subclone size inferred using two-way cluster modeling, we can determine the order of any pairs of recurrent mutations existing in multiple samples. Herein, we use the notation MCP 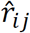 to indicate the estimated MCP of mutation *i* from the model of sample *j*. If a pair of mutations is recurrent in tumors with a fixed order, the relative size of their estimated MCPs should be consistent. For any two loci *a* and *b* with somatic mutations, the MCP profiles across p samples are 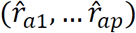 and 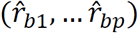. To determine whether the two mutations are highly related, the Wilcoxon signed-rank test can be applied to the profiles of the two mutations. In the event of significant inequality, when one mutation is more common in cells than the other mutation, it indicates a recurrent evolutionary order between the two mutations.

## Results

In this study, we first evaluated the prediction accuracy of CloneDeMix by simulated data. Simulation study is useful to verify how well an algorithm behaves with data generated from the theory, but it cannot inform us how well the theory fits reality. To that end, we collected normal RNA sequences from TCGA and applied down-sampling to these normal data to create artificial copy number changes. We used the data to compare CloneDeMix with THetA by evaluating weighted root mean square error of MCP estimation and positive rate of copy number prediction. We also applied CloneDeMix on head and neck cancer data from TCGA and serial biopsies of esophageal cancer [19] to infer genomic evolution based on copy number change.

### Simulation

The simulation considered four variant states of copy numbers, namely 0, 1, 3, and 4 copies. Four MCPs were included: 0.1, 0.3, 0.5, 0.7, and 0.9. Each combination was repeated three times, thus resulting in 60 regions with copy number changes. Furthermore, each region was assigned 20 bases generated with a Poisson distribution whose mean value was determined by its assigned copy number state and MCP. In addition to the mutated regions, 100 normal regions were scattered among the mutated regions; their copy number state was two. The simulation generated depths for a long sequence with 3,200 bases for each of the 60 samples. CloneDeMix was subsequently applied to the simulation data to reconstruct respective copy number states and MCP groups. The entire simulation process was repeated 10,000 times to obtain a conclusion.

The simulation was performed to evaluate the model estimation accuracy. Table 1 shows the mean and standard deviation (SD) of simulation results for MCP estimation, and Fig 4 demonstrates the accuracy of assignments for copy number states. As presented in Table 1, the MCP estimates were very close to the underlying truth, indicating high performance for MCP estimation. Notably, the bias decreased as the ground value of the MCP increased. Detecting mutations of low cellular prevalence was relatively difficult because the signal was not adequately strong.

**Fig 4.**
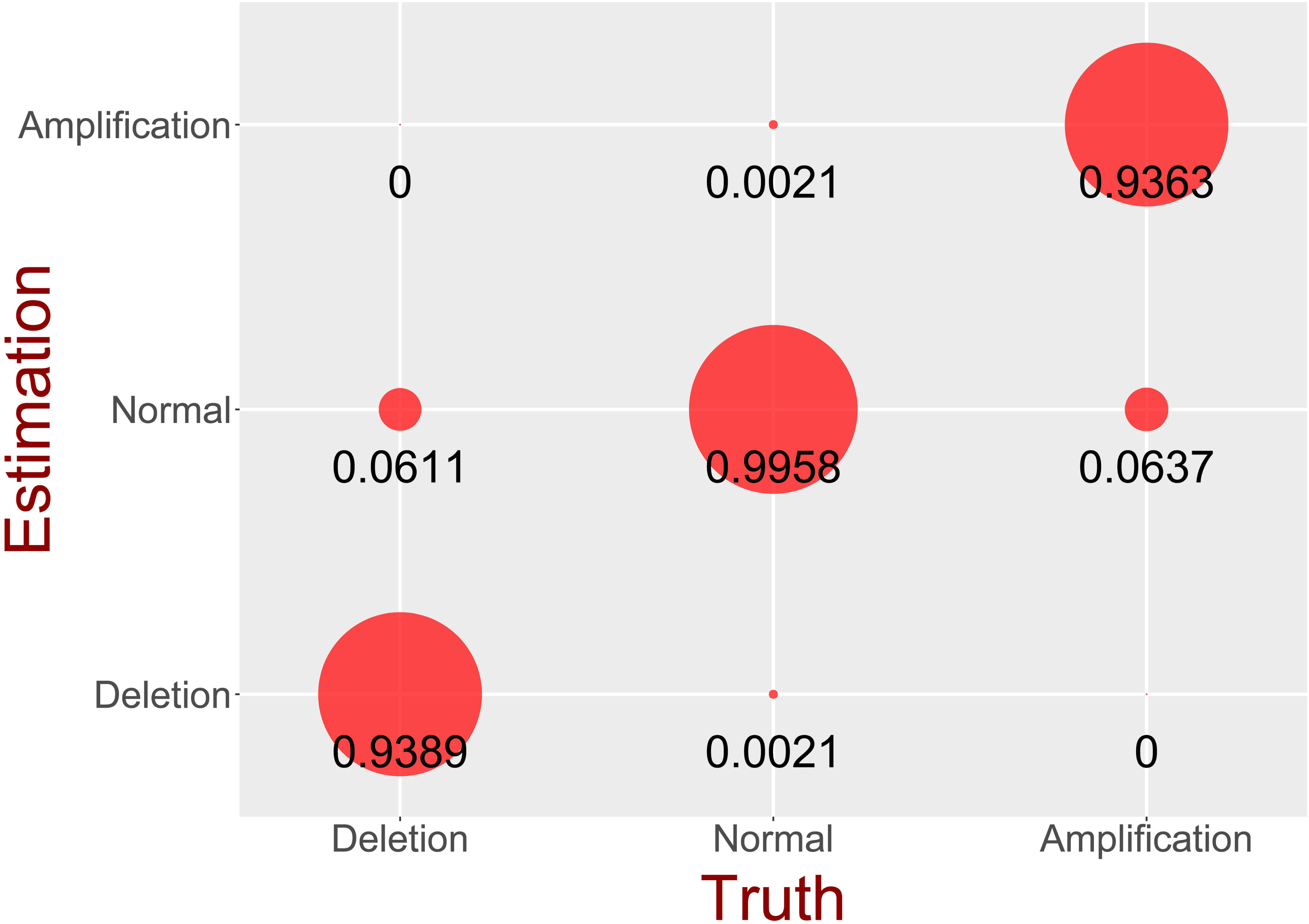
Result of copy number estimation. The size of the circle is proportional to the number of loci assigned to each estimated status from 10,000 simulations. The CNA status is divided into three conditions: deletion, amplification, and normal conditions.

**Table 1.**
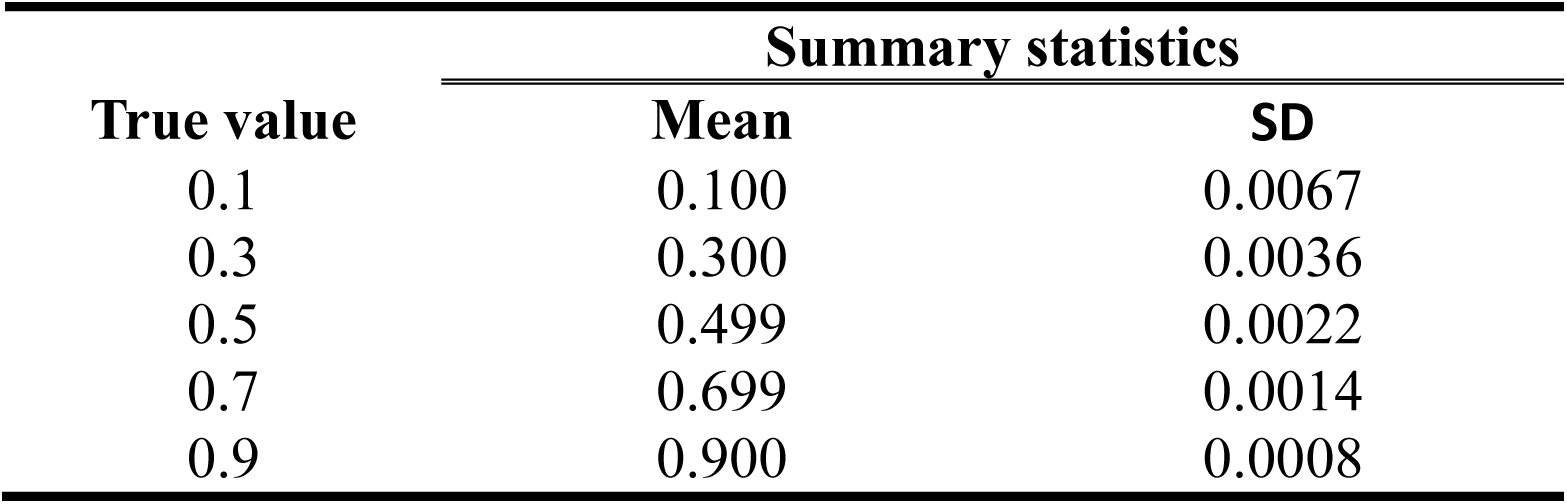
Mean and SD of MCP estimation.

| True value | Summary statistics |  |
| --- | --- | --- |
|  | Mean | SD |
| 0.1 | 0.100 | 0.0067 |
| 0.3 | 0.300 | 0.0036 |
| 0.5 | 0.499 | 0.0022 |
| 0.7 | 0.699 | 0.0014 |
| 0.9 | 0.900 | 0.0008 |

As illustrated in Fig 4, the accuracy of the copy number assignment under each condition was calculated from 10,000 simulations. The specificity of CloneDeMix was found to be 99.58%, and the sensitivity for amplification and deletion were 93.65% and 93.89%, respectively. Thus, CloneDeMix represents a high specificity and efficiently controls false positive results. As mentioned in the discussion on MCP estimation, estimating mutations of low cellular prevalence was biased. These biased MCPs directly caused the misclassification of the copy number state and reduced the model sensitivity. In conclusion, these simulations support that CloneDeMix can successfully identify the potential copy number mutations and deconvolute its amplification or deletion state from the clonal architecture.

### Comparison with THetA2

In this section, we evaluated CloneDeMix on a more realistic simulation scenario and compared it with THetA2. The core concept of this simulation scenario is the use of down-sampling technique to resample reads of real normal sequencing data with artificial copy number changes.

To that end, we first collected 75 normal samples from TCGA and then performed standard quintile normalization to reduce noise. For simplicity, we only used chromosome 1 for validation, and chromosome 1 was first cut into 200 different regions. According to the raw data, we have the raw read counts of each region per sample. The 75 samples were equally divided into case and control. In the control group, the resampled read count of each region was generated from a binomial distribution. For the parameter setting of a binomial distribution, the number of trials is set as two times raw read count and the success probability is 0.5. This procedure is called down-sampling and it guarantees the mean of resampled count is the same as the mean of raw count. In the case group, we need to randomly assign 20 regions to have copy number change. The resampled read count of CNA region also followed the binomial distribution with the number of trials equal to two times of the raw read count, but the success probability is set as 0.5×(2×(1-MCP)+C×MCP)/2 which is determined by a predefined copy number C and MCP. The predefined copy number of a variant was set to be 0, 1, 3, and 4. The MCP was set to be 15 different values ranging from 0.1 to 0.9 as shown in Fig 5.

**Fig 5.**
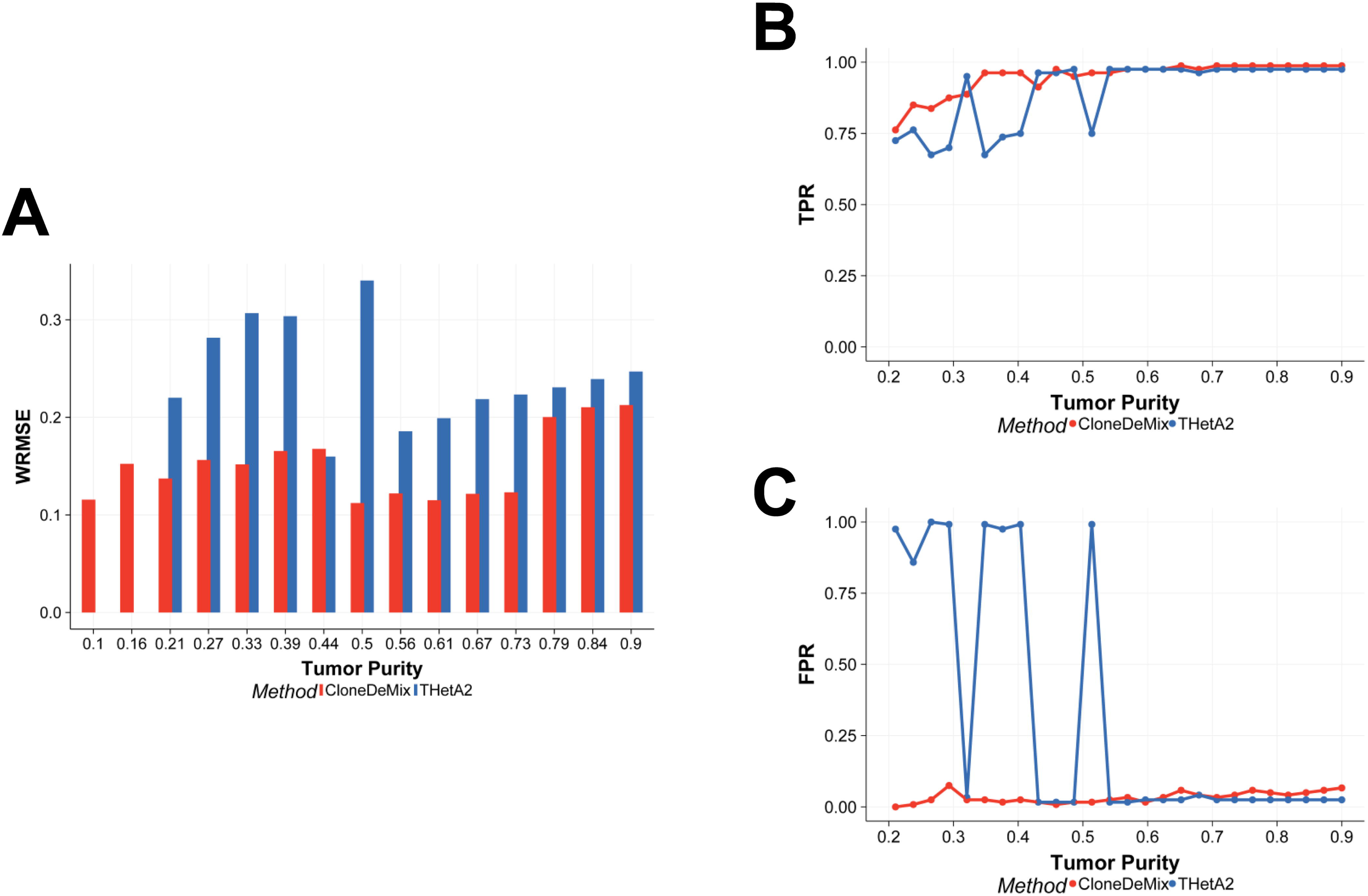
Comparison of CloneDeMix and THetA2 with resampled data. (A) The Y-axis is the weighted root mean square error (WRMSE) for measuring the performance of MCP (or purity) estimate, and X-axis represents the true purity setting. (B) The true positive rate (TPR) of copy number detection. (C) The false positive rate of copy number detection.

Most studies integrate CNAs and single nucleotide change to improve the accuracy of copy number identification and to reduce the bias of cellular prevalence estimation. However, those approaches only study the regions that contain single nucleotide change, and this constraint apparently limits our understanding of the chromosome structure change. It has been reported that CNAs affect a larger fraction of the genome in cancers than any other type of somatic genetic mutation does [23]. For example, a large-scale study of somatic CNAs across different cancers shows that in a typical cancer sample, 17% of the genome was amplified and 16% genome was deleted on average [24]. Hence, for a fair comparison, we only compared CloneDeMix with THetA2 because THetA2 is also a subclonal copy number decomposition method and supports direct tumor heterogeneity inference without considering SNVs.

Both of CloneDeMix and THetA2 are developed for multiple clone identification, but THetA2 tends to identify single clone in our experience. Therefore, we designed the resampled data as a mixture of normal cells and one subclone of tumor cells. In this simple case, the MCP is equal to the tumor purity and we explored the model performance in different purity. In Fig 5A, we measured the performance of purity estimation by weighted root mean square error (WRMSE) which is a type of adjusted RMSE. WRMSE adopts the inverse of true purity as the weight for adjustment because the variance of purity estimation is a function of the true purity. The variation of purity estimate increases when the purity increases. Across the 15 different purity settings, CloneDeMix outperforms THetA2 on measuring purity as demonstrated in Fig 5A. It is notable that the WRMSEs of THetA2 are missing zero in Fig 5 at low purity settings (0.1, and 0.16) because THetA2 cannot identify tumor population at low tumor purity. We calculated the true positive rate (TPR) and false positive rate (FPR) of copy number assignment at different purity levels in Fig 5B and Fig 5C. We found that both of them performed well when tumor purity was larger than 0.5. CloneDeMix outperformed THetA2 in the low purity. It indicates CloneDeMix and THetA2 are equally well at exploring large subclones while CloneDeMix has better detection power for small subclones.

### Preprocessing of TCGA data

We analyzed the whole-exon sequencing data of 75 head and neck tumor samples with their paired normal samples from TCGA (http://cancergenome.nih.gov/). This dataset includes a total of 20,846 genes with 180,243 exons. We assumed the copy number state of a single exon to be homogeneous. Each exon was represented by the mean read depth. The read-depth profile of a tumor sample was normalized with loess transformation against its paired normal sample. The baseline parameter *a_base,i_* for exon *i* was estimated from the paired normal sample by using half of the read depth of the normal sample at the same locus. Because the normal sample could also have an abnormal copy number status, we checked it against all other normal samples. The target normal sample was first normalized against all other normal samples by using the cyclic loess method and was subsequently processed through CloneDeMix to identify the copy number status at each locus. In this step, the average profile of all other normal samples was treated as the baseline. If, for example, an abnormal copy number is found to be k, the raw read depth of this locus would be divided by k to provide the estimate of *a_base,i_* for tumor modeling.

### Copy number distribution and clone structure

We applied CloneDeMix to each normalized sample and estimated the copy number state of each locus as well as the corresponding MCPs. Fig 6A shows the chromosomes that were mutated most frequently, and the results of all other chromosomes are shown in S1 Fig. This figure presents the copy number events across 180,243 exons for each of the 75 tumor–control sample pair. The proportion of exons with a normal copy number was high in all samples, and it was close to 100% in the control samples. The proportion was significantly decreased in the tumor samples, indicating considerable structural variations during cancer development.

**Fig 6.**
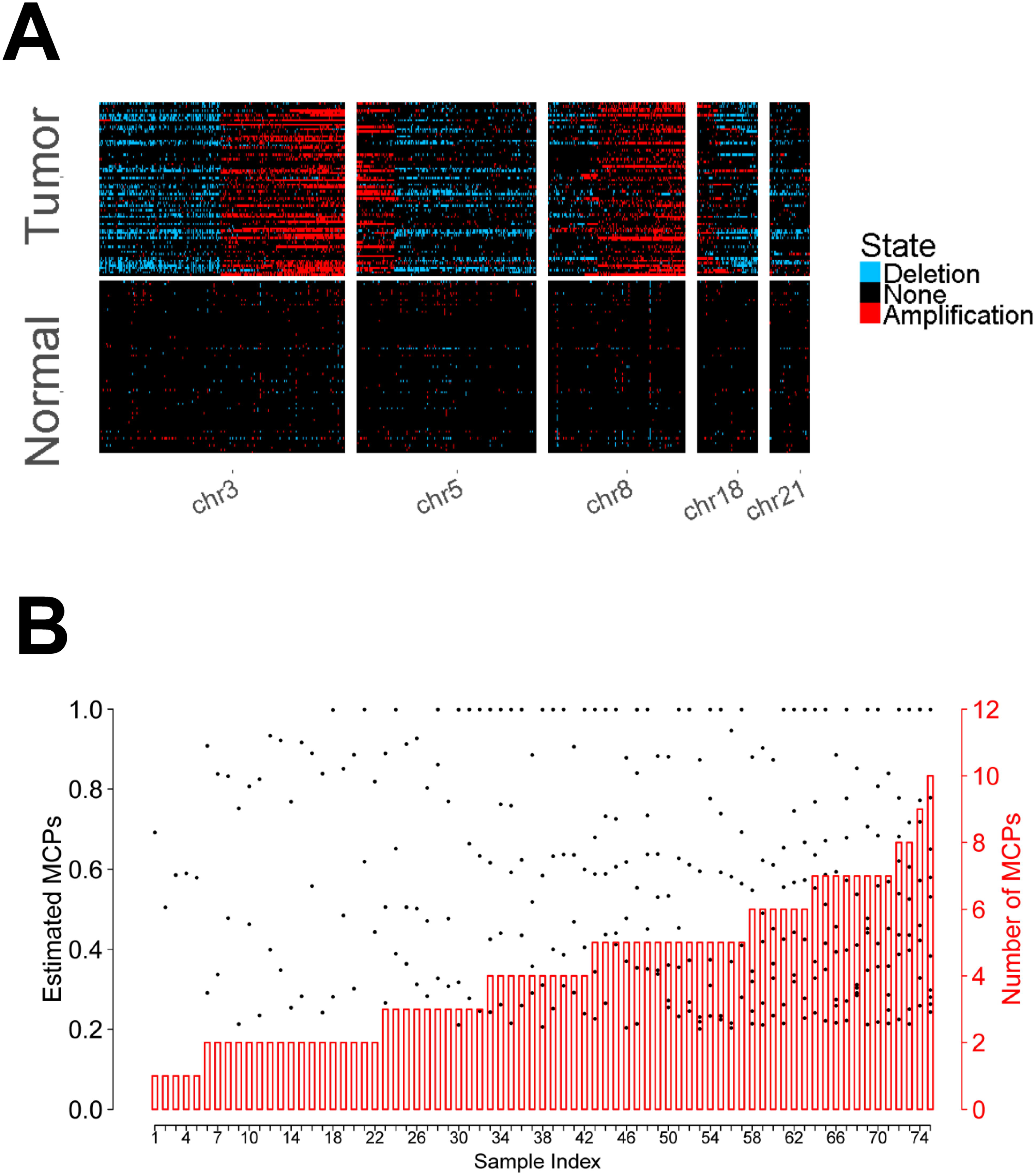
Copy number estimation of chromosomes with high mutation rates. (A) The estimated copy number states for exons across the genome are presented by different colors. Light blue and red represent the deletion and amplification events, respectively. Black indicates no copy number changes. (B) The black dots indicate the estimated MCPs with respect to the left axis. The red bars represent the number of MCPs with respect to the right axis.

On average, 4.7% and 8.7% of exons were estimated to have deletion and amplification, respectively. We also found that the exons located at 3p, 21p, and 18q were deleted most frequently, and the average proportions of deletion within these chromosomal arms were 19%, 17%, and 13%, respectively. Conversely, the estimated amplification frequently occurred at 3q, 8q, and 5p, with average frequency levels of 29%, 24%, and 23%, respectively. Previous studies have reported a loss of 3p and 8p as well as gains of 3q, 5p, and 8q not only in head and neck cancer but also in most tumors [20-23]; these results are concordant with our findings. Other novel subclonal CNA regions that were not reported in pan-cancer data analysis [20-23] were identified as multiple tumor subpopulations were considered (e.g. Deletion in 21p, S1 Fig). These subclonal CNA signals may be diluted in the previous studies that assumed only one homogeneous tumor clone and inferred CNAs from the average of whole tumor information. Our results confirm the identification strength of large-scale structural variations based on clonal evolution.

Fig 6B presents a summary of MCP estimation. The number of MCPs was determined using the model selection criterion AIC. We associated the number of subclones in the tumors with clinical outcomes because this number is closely related to tumor heterogeneity. The target phenotype included tumor invasion and metastasis, which are particularly ominous signs of poor prognosis in head and neck cancer. The association analysis was applied to only 68 samples because the clinical records of the other samples were incomplete in TCGA. Fig 7A illustrates the box plot of the number of MCP groups under each clinical group. In this figure, a sample is denoted as “NO” if no record of either invasion or metastasis exists; otherwise, it is denoted as “YES.” There appeared to be a tendency of increased tumor heterogeneity for tumors with invasion or metastasis. The variation of numbers of MCPs was larger for this group. To more comprehensively clarify this factor, we dichotomized the number of MCPs into two groups. The number of MCPs exceeding 4 indicated strong tumor heterogeneity, whereas a lower number indicated less heterogeneity. The contingency table (Fig 7B) shows the dichotomization of tumor heterogeneity associated with the clinical outcomes. The corresponding odds ratio was 3.64, and the p value evaluated with logistic regression was 0.029. For the samples with higher tumor heterogeneity, the odds of invasion and metastasis were 3.64 times higher than those for the samples with lower tumor heterogeneity. In recent studies of head and neck cancer, this association between tumor heterogeneity and metastasis was explored by whole exome sequencing and single cell RNA sequencing [25-27]. These studies also found the difference in tumor heterogeneity between primary and matched lymph node metastases samples.

**Fig 7.**
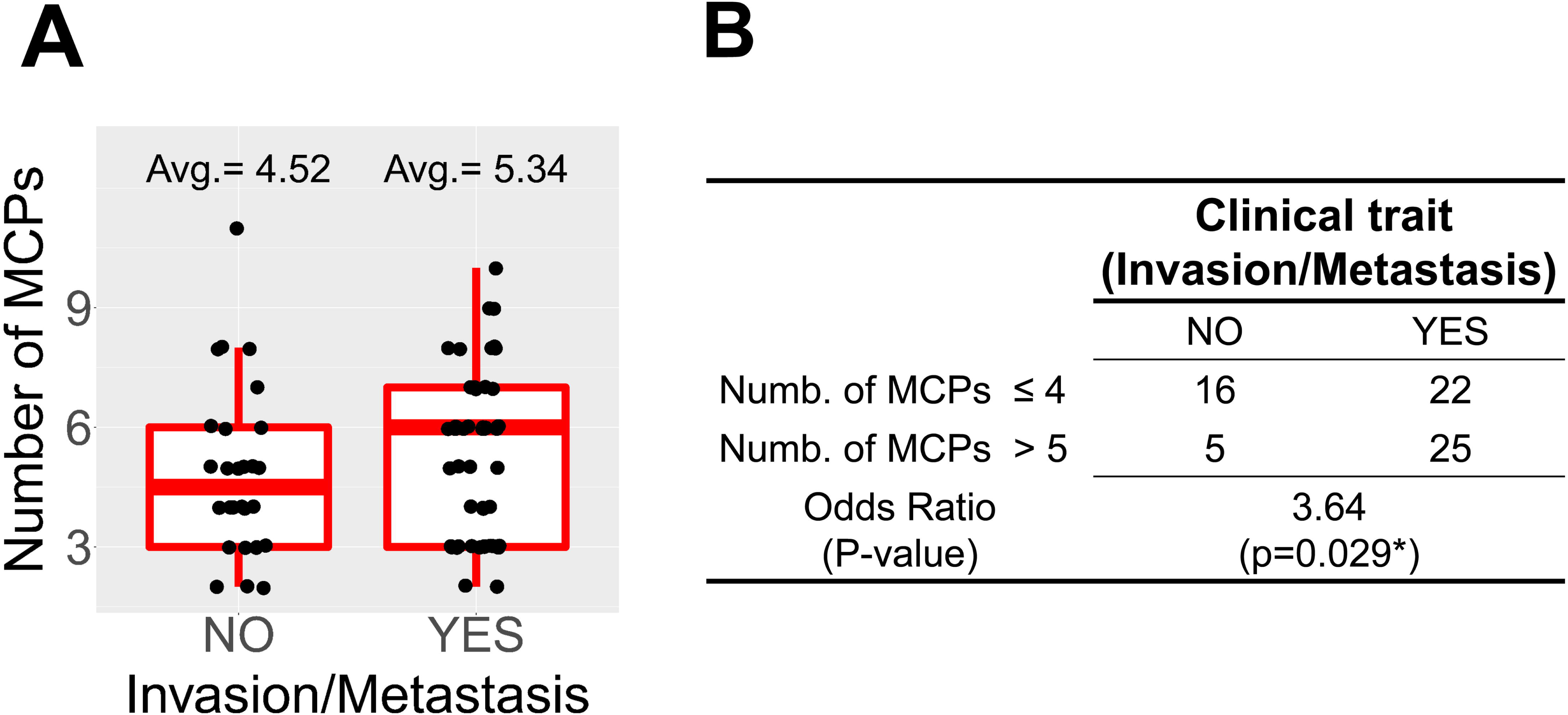
Comparison for the number of MCPs in different clinical groups. (A) The box plot for the number of MCPs with and without invasion or metastasis. The number of MCPs in each sample is represented by a black point jittered around the box. (B) Contingency table for dichotomization of tumor heterogeneity and clinical outcomes.

We further investigated the association of overall patient survival and tumor heterogeneity by survival analysis, and used two different ways to demonstrate this association. First, we directly considered the subclone number as a covariate of survival analysis, and then applied Cox model to analyze the effect of subclones. We got a p-value, 0.036, by Wald’s test, and apparently tumor heterogeneity is a risk factor for survival. Next, we considered three different tumor heterogeneity levels of samples and performed Kaplan-Meier (KM) curve for different levels. To this end, all of the samples are divided into three classes by its subclones number, low-heterogeneity (less than 5 subclones), median-heterogeneity (5 ≤ subclone number ≤ 8), and high-heterogeneity (large than 8 subclones). The sample sizes of the three classes are 20, 36, and 19, respectively. Fig 8 showed the survival curves of the three classes with different colors, and the survival curve of high-heterogeneity samples is worse than the others. Hence, high-heterogeneity is associated with poor overall survival. It indicates the tumor behavior varies with its heterogeneity. The heterogeneity and mortality in head and neck cancer was also investigated by a different approach [26], and it also concluded that high-heterogeneity in tumors had doubled the hazard of death.

**Fig 8.**
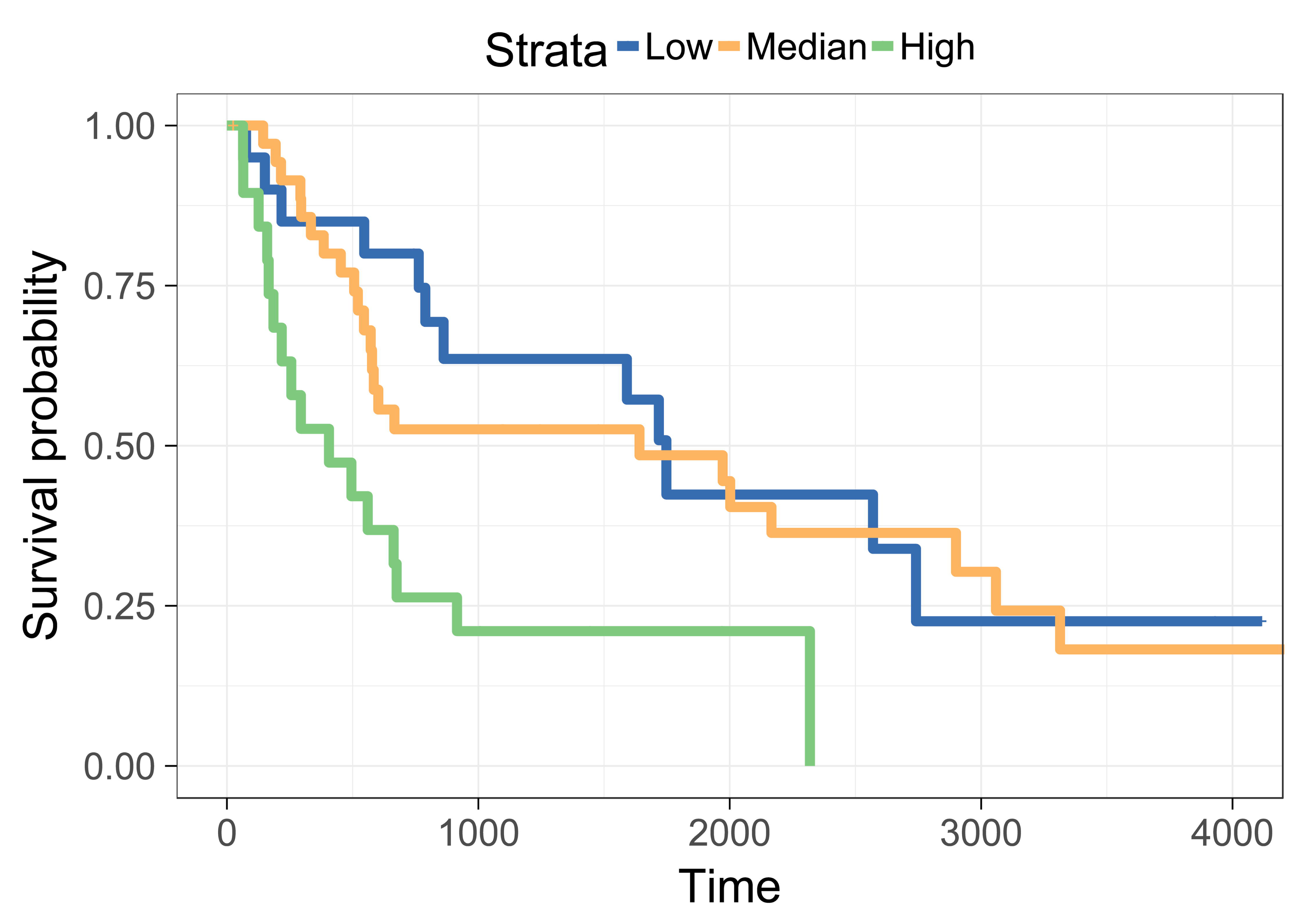
Survival curves between different classes of heterogeneity levels. There are three Kaplan-Meier (KM) curves. The blue, yellow, and green represent the group of low, median, and high heterogeneity, respectively.

### Inference of evolutionary order of mutations

As stated in the Methods section, we inferred the evolutionary order of recurrent variants with multiple samples. For easy comprehension, we demonstrated the result at the gene level through a series of summary steps. We first selected the genes with consistent amplification or deletion states in more than 25% of the exons within at least one sample. A total of 3,244 genes were included in this demonstration, and this set is called the background gene set. For each sample, the MCP of a gene was represented by the mean MCP of its exons. We then performed the Wilcoxon signed-rank test using the gene-level MCP of any two genes across the samples to derive all pairwise evolutionary relationships. For example, if the MCP of gene *i* was larger than that of gene *j* (p = 0.05), the mutation on gene *i* was more likely to be an earlier event than that on gene *j*. This relationship was marked as 1; otherwise, it was marked as 0. The 0–1 matrices of pairwise evolutionary relationships were separately calculated for samples with and without nodal metastasis, and they could be denoted as a matrix *M_neg_* and *M_pos_*. The element of the matrix could be denoted as *M_E,ij_*, representing the evolutionary order of mutations on gene *i* against mutations on gene *j* inferred with samples under the E condition, which could be *neg* or *pos*.

The evolutionary order matrix can be used to construct an evolutionary tree of all mutations. However, a tree of 3,244 genes is highly complicated, rendering the comparison of different clinical traits difficult. Therefore, for simplification, we proposed a progression score to summarize the relative position of a mutation on the evolutionary tree of tumor formation. The scores of a gene in advanced tumors can be compared with those of genes in newly developed tumors to select the ones that recurrently occur in the early stage of tumor development. The P score of gene *i* under condition E is thus defined as a summary statistic from the evolution matrix and is formulated as follows:

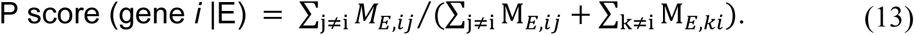

Among all relations of gene *i* with other genes, the P score provides the number of times the mutation in gene *i* is more likely to occur before that in other genes. If a gene is close to the root of an evolutionary tree, its corresponding P score must be higher than that of its descending gene.

We first investigated the P-score behavior of prevalent genes which have been discussed in head and neck cancer [22], and the results are listed in Table 2. The P-score of PIK3CA is consistently larger than 0.9 across different clinical traits. That is, the mutation of PIK3CA occurs early in the tumor progression. In contrast, patients with perineural invasion acquire early mutation of CDKN2A gene more often. Some of the well-known cancer genes are not powerful in our P-score analysis. For example, we identified structure variation of TP53 only in a few patients, and these few MCPs are not enough to construct a powerful P-score.

**Table 2.**
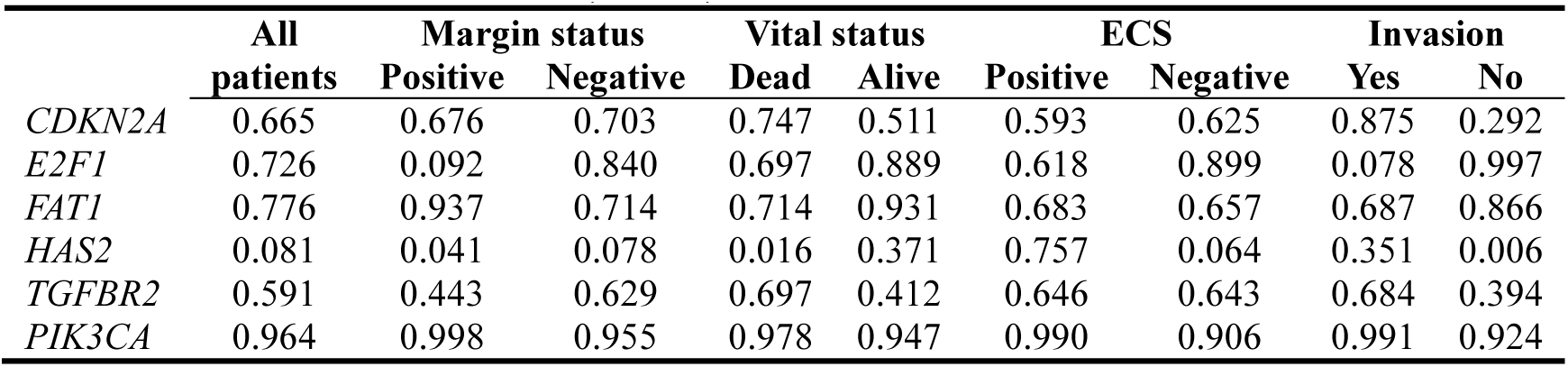
The P-score of CDKN2A, E2F1, and PIK3CA in different clinical outcomes.

We compared the P score between the samples with and without nodal metastasis by plotting a scatter plot (Fig 9). Most background genes tend to mutate in a random order not related to tumor progression. According to our P score definition, we postulated that the driving genes of lymph node metastasis would be scattered above the diagonal line. The genes above the diagonal line of the plot are more likely to acquire mutations at an earlier stage of tumor formation and occupy a significant proportion of the tumor at its advanced stage. This would yield higher P scores when only the samples with lymph node metastasis are considered. By contrast, the prevalence of mutations in those genes might be low in the samples without lymph node metastasis and hence yield lower P scores.

**Fig 9.**
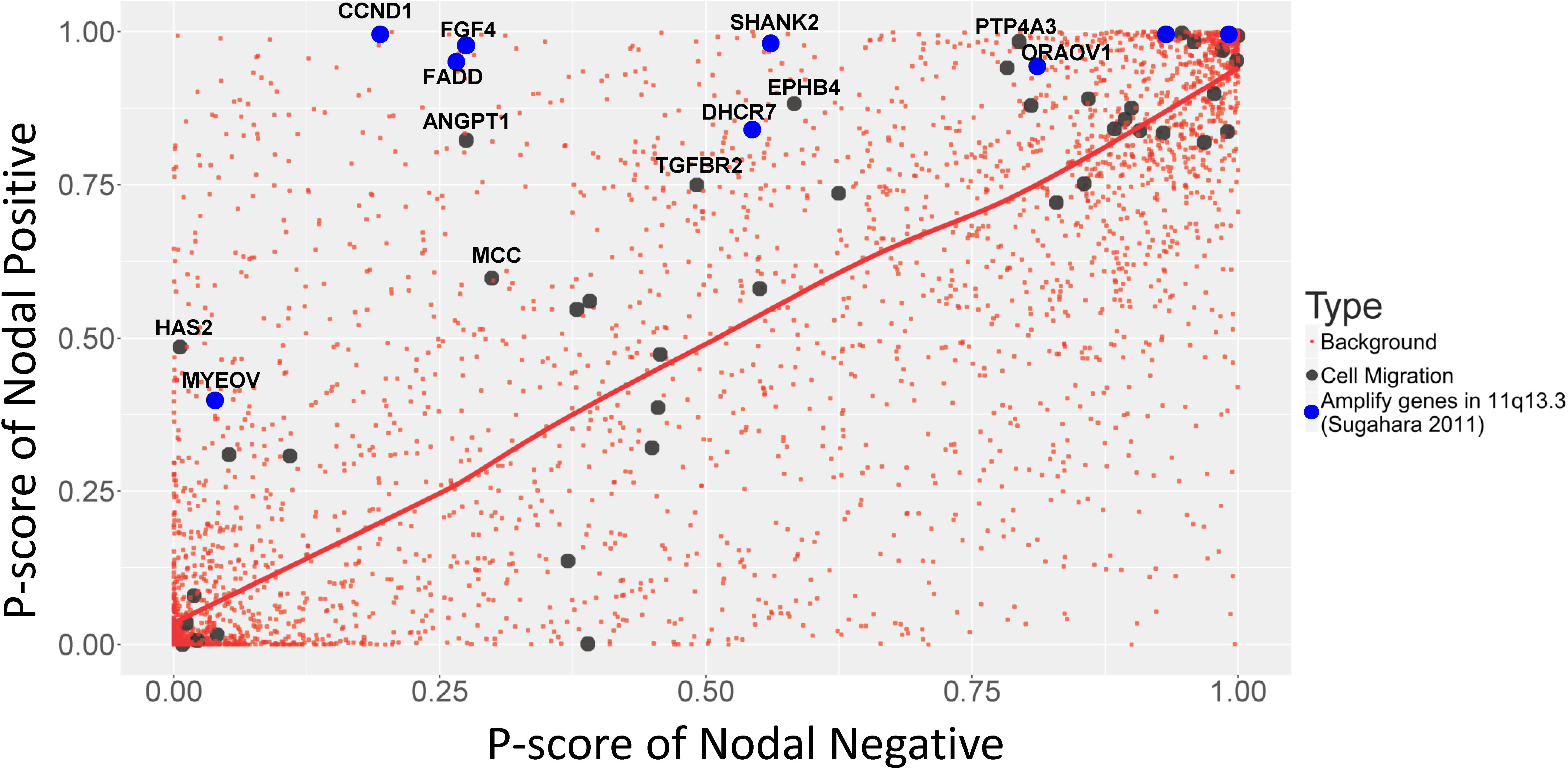
Scatter plot of P scores between nodal positive and nodal negative samples. The red points indicate the background gene set. The red curve indicates the loess smoothing curve constructed using all points in the figure. Genes related to cell migration are marked in black. The genes from 11q13.3 are marked in blue. The literature supporting genes are labeled.

To confirm our conjecture, we selected the genes by their biological functions using ConsensusPathDB web (http://cpdb.molgen.mpg.de/) and investigated whether genes related to metastasis in the literature are more likely to be distributed above the diagonal line. Because cell migration is a crucial step in the metastatic cascade, we selected cell-migration-related genes, which are marked as black in Fig 9. Consequently, we found that 43 genes had the function of cell migration. Most of these genes were distributed above the diagonal line of the P score scatter plot, whereas some were distributed below the diagonal line. Recurrent mutations in these cell migration genes are expected to be the driving forces for the initiation of lymph node metastasis, consistent with our observations. For example, HAS2 is a member of the gene family encoding putative hyaluronan synthases, which control the biosynthesis of hyaluronan and critically modulate the tumor microenvironment. Several studies have shown that the inhibition of HAS2 reduced the invasion of oral squamous cell carcinoma [28-30]. Similar to HAS2, ANGPT1 is located in the upper left corner and has been recently investigated for the mechanism of lymph node metastasis [31-34]. ANGPT1 plays an important role in the regulation of vascular development and maintenance of vessel integrity. A study showed that the activity of ANGPT1 induced the enlargement of tumor blood vessels to facilitate tumor cell dissemination and increased the ability of metastasis in tumors [34]. Fibroblast growth factor (FGF)-4 is another notable example. The P score of FGF4 significantly differs in nodal positive and negative patients. FGF4 is a member of the FGF family and possesses broad mitogenic and cell survival activities. It has been proposed to be involved in tumor growth, cell proliferation, and lymph node metastasis [35-37]. In contrast to the black genes located in the upper left corner of the plot in Fig 9, few studies have reported any relationship between the black genes located in the lower right corner and lymph node metastasis, although they have the same biological function. A complete literature review of the genes associated with cell migration and tumor metastasis is presented in S1 Table. The observations suggest that our inference of the clonal evolutionary order is relevant and can be applied for identifying causal drivers.

Another notable observation is about the neighboring genes of FGF4. As mentioned, FGF4 is an important gene for driving lymph node metastasis. It is located in 11q13.3, which is frequently amplified in head and neck squamous cell carcinoma [35]. Sugahara also listed several other genes in 11q13.3 that are related to cancer development, namely *TPCN2*, *MYEOV, CCND1, ORAOV1, TMEM16A*, *FADD, PPFIA1, CTTN, SHANK2*, and *DHCR7*. We also assessed their status by using the P score analysis; the genes are indicated in blue in Fig 9. All these genes were above the diagonal line. Their corresponding P scores showed considerably significant differences between patients with and without nodal metastasis. Hence, we postulated that those genes in 11q13.3 are possibly related to lymph node metastasis in head and neck cancer. Several previous studies have confirmed this observation, as reported in S2 Table.

### Application on serial biopsies of esophageal cancer

We next applied CloneDeMix on multiregional whole-exome sequencing data from 13 primary esophageal squamous cell carcinoma (ESCC) patients [19]. There are 51 tumor regions and 13 matched morphologically normal esophageal tissues sequenced with the mean coverage of 150×. For fair comparison, we selected 11 of 13 patients based on its platform. We also removed patient ESCC07 because we only got two regions successfully aligned to the reference genome. In total, we included 10 patients in this application, and, for each patient, we have four different tumor regions with one matched esophageal tissue. As preprocess of TCGA data, the read-depth profiles of ESCC tumors are normalized with loess transformation against its paired normal sample. For each individual, the paired normal tissue is also used to calculate the estimates of baseline, and then applied CloneDeMix to tumors for gene-specific CNVs and MCPs.

In this application, we aim to explore the variability of evolutionary structure among multiregional tumors by inferring the order of copy number change. For the purpose of studying variability between regions, we only focused on the frequently mutated genes which are informative about tumor evolution. Although the construction through these genes is not able to resolve completely the entire evolutionary structure, the inferred structure between regions can still facilitate the understanding of tumor progression. To that end, we collected the target gene list from the Ion AmpliSeq Comprehensive Cancer Panel which includes 7,044 exons of 409 tumor suppressor genes and oncogenes. The estimated CNVs and MCPs of the ESCC biopsies for this gene set were summarized and interpreted as follows.

We first investigated genomic heterogeneity of ECSS through MCP comparison. MCP is a gene-specific measurement of fraction of cells that carry a certain mutation, and we can study the overall structure of MCPs across whole genome to reveal the genomic heterogeneity of a given sample. We calculated the correlation matrix of MCP between samples, and this correlation matrix is presented in Fig 10. The diagonal blocks of this correlation matrix are tissues of the same sample and are slightly higher than the others. The average correlation of diagonal block is 0.5 and the average of off-diagonal cells 0.3. It shows that the MCP structure within each patient is more consistent than between patients.

**Fig 10.**
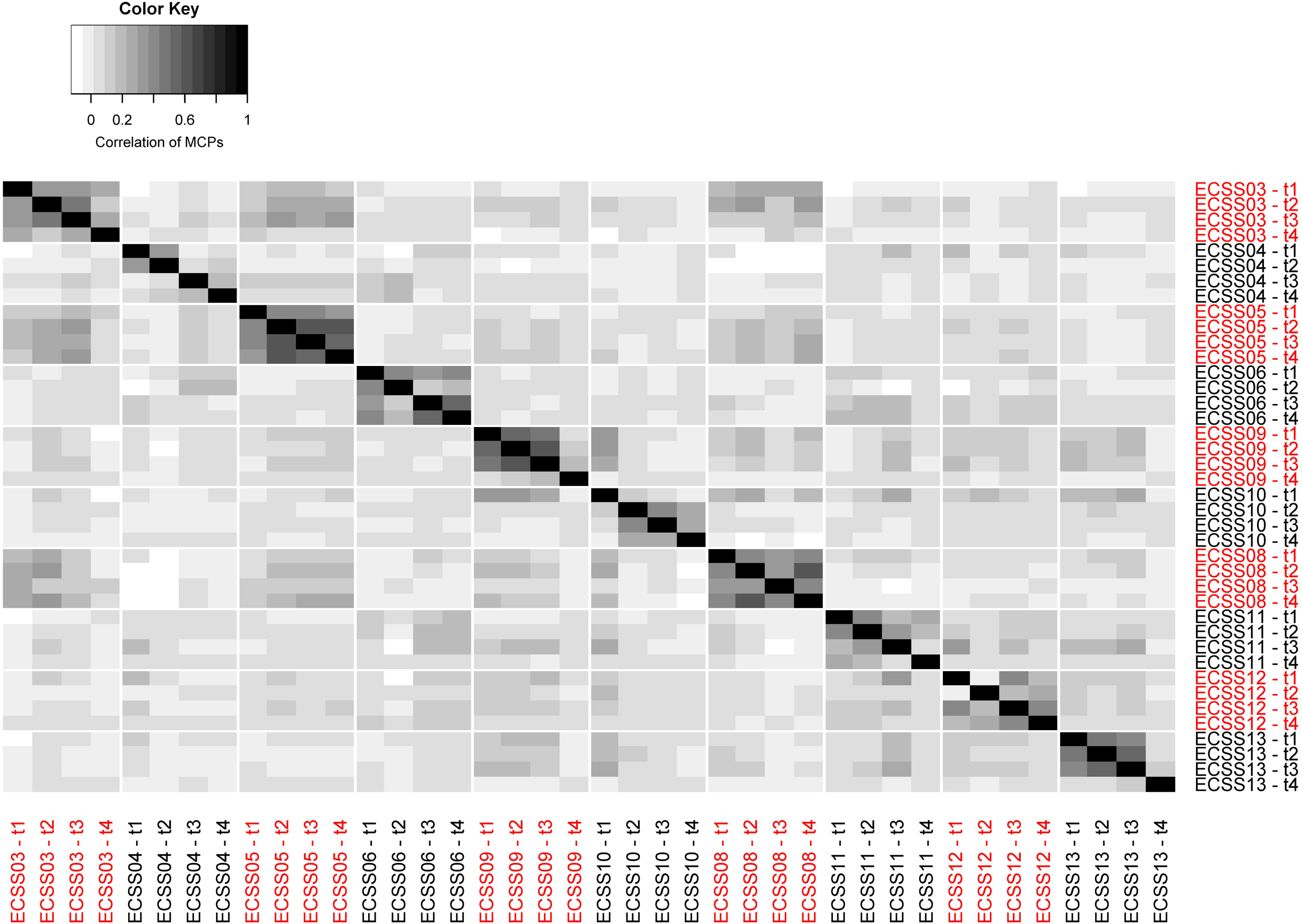
Correlation matrix of MCP between samples. Each cell indicates the correlation of MCPs between the corresponding ESCC samples.

Next, we identified the evolution-related genes for each individual. In ESCC study, each tumor was dissected into four regions, and this kind of serial biopsies has a natural assumption that the size of MCPs is comparable within a given tumor. This characteristic can facilitate the individual-specific heterogeneity study. In order to explore individual-specific heterogeneity, we first identified genes on the trunk and on the branch of the evolutionary tree separately. The trunk of the tree refers to the CNAs consisted in all regions while the branch refers to those only in some regions. We can identify these genes according to the MCP across regions. A gene is located on the trunk of a tree if its average MCP across four regions is larger than 0.8, and a gene is located on the branch of a tree if the MCP of one region is larger than the average MCP of all the remaining regions by 0.7. Instead of tree comparison, we directly compare the MCP matrix of selected genes (Fig 11). In Fig 11, genes in red rectangles are selected to be the trunk genes, and the remaining genes are on the branch of a tree.

**Fig 11.**
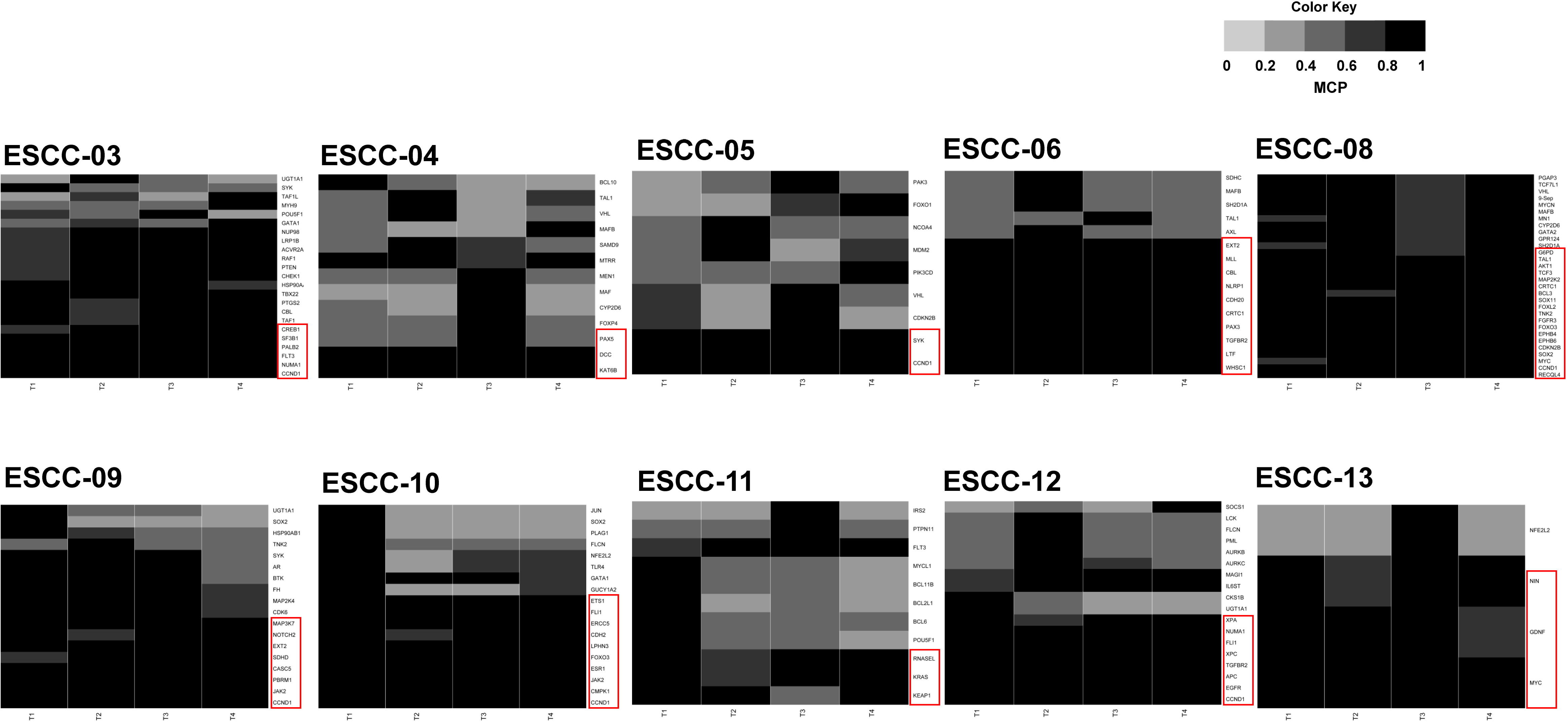
MCP matrices of selected genes among 10 samples. There are six MCP matrices. The color of each cell represents the MCP quantity of a gene for a given sample. The labels of rows indicate the gene symbols, and the labels of columns are region index A gene within the red rectangle is identified as the gene located on the trunk of an evolutionary tree.

The two types of genes defined above reveals huge variability of evolution structure across tumors. The genes on the trunk of a given tree represent the genes changed in copy number at an earlier stage of tumor formation, and these genes have potential ability to drive tumor growth. For example, most of the genes on the trunk of sample ESCC12 (CCND1, EGFR, APC, TGFBR2, XPC, XPA, FLI1, and NUMA1) have been identified and initially reported on the esophageal cancer [38-42]. Although the genes on the trunks of trees vary among different individuals, there are still genes repeatedly identified in multiple individuals such as CCND1, JAK2, UGT1A1, FLI1, NFE2L2, SOX2, CDKN2B, and MYC. Specifically, CCND1 was identified in six individuals as the trunk gene and is a well-known cancer oncogene located on 11q13. Its amplification has been reported in several human neoplasias [43].

## Discussion

In this study, we developed CloneDeMix for the deconvolution of tumor progression through high-throughput DNA sequencing data. The features of CloneDeMix are as follows. First, it reconstructs an evolutionary structure of copy number changes during tumorigenesis. Most existing methods for cancer evolution discuss the history of single-nucleotide changes and derive the potential driver genes. However, the importance of CNAs is growing and its influence on disease and cancer development is clearly established [44]. Therefore, the reconstruction of copy number evolution in tumor progression is in demand. Second, CloneDeMix provides the MCP as a measure of the evolutionary structure. This measurement is used to estimate the fraction of cells containing a specific set of mutational events. According to the definition of the MCP, it provides a more direct evolutionary reconstruction than does the SCP, which is defined as the size of a subclone in a tumor. For instance, the MCPs of early mutations in cancer must exceed those of other mutations, but no such structural relationship exists for SCPs. Although MCPs of a tumor is related to its phylogenetic tree, we do not have DNA haplotypes to resolve the tree architecture from many possibilities for each individual tumor. Hence, in this study, we only borrow the strength of multiple samples to understand potential evolutionary orders using the P score. Third, our model exhibits high flexibility. CloneDeMix can identify the copy number state of any type of variant, from a single nucleotide to a moderate size of regions. Furthermore, the model facilitates the simultaneous analysis of multiple types of targets because it depends on only the summary information of each locus.

The simulation study revealed that CloneDeMix can identify the current clonal structures of a tumor. The accuracy of copy number states was nearly 93%, and the MCP was also accurately restored (Table 1). Furthermore, the application of CloneDeMix to head and neck cancer data from TCGA yielded promising putative CNAs. The deletions observed on chromosomes 3p, 18q, and 21p and the amplifications on chromosomes 3q, 5p, and 8q are consistent with most cancer studies on copy number identification [20-23]. This observation strongly supports our CNA inference procedure.

When the estimation accuracy reaches a certain level, the most important concern is to understand the relationship between tumor heterogeneity and disease progression. Tumor clone dynamics have been associated with clinical outcomes for different types of cancer [45-47]. Our method provides a quantitative measurement of clonality, and it is associated with tumor invasion and metastasis development in TCGA database. Tumors with more subclones are a result of complex branched evolution, implying a series of adaptations to a new environment. These newly emerged subclones may contribute to metastatic initiation or acquire a new ability to invade the lymphatic or vascular system. Thus, the strong prognostic association of the number of MCPs with invasion or metastasis reinforces its clinical relevance; this index appears to be a novel feature for further exploration.

We established a novel score, the P score, for evaluating the order of a recurrent mutation in the evolutionary hierarchy by analyzing multiple samples. By comparing the P scores of a somatic variant between different clinical groups, we could identify the copy number mutations that occur early in the tumor stage and expand the accompanied subclones with time. The utility of P scores was also demonstrated in the head and neck cancer data according to the sample status of metastasis. Furthermore, we identified a group of genes that matched this condition. Specifically, the genes located at 11q13.3 are well known to be frequently amplified in head and neck squamous cell carcinomas. Their P scores in our analysis were particularly high for the samples with lymph node metastasis and relatively low for those without metastasis. Accordingly, those gene amplifications are potential causal mutations to drive metastatic cascade in head and neck cancer. Hence, screening for genes that differ considerably in their P scores is meaningful for driver gene detection.

The success of our approach highly depends on the coverage of DNA sequencing. A higher read depth can more efficiently reflect the clonal structure and copy number changes of different loci. Currently, CloneDeMix makes an independent assumption without considering the dependency among closely located loci. Hence, the neighboring segments are not grouped into the same copy number events. This can be an advantage as well as a disadvantage because there is no clear understanding about the range covered by a copy number event. Technically, we can still integrate the correlation structure into CloneDeMix to improve the flexibility; this is an ongoing project for our next version of the R package.

CloneDeMix can easily integrate different types of somatic mutations detected in sequencing data. For example, the well-studied SNVs carry extensive information about the clonal expansion in tumors. CloneDeMix can consider the copy number status of two alleles individually if the detection of each allele is optimized. Therefore, we expect CloneDeMix to be useful in understanding tumor heterogeneity and how it evolves to the current status. Moreover, CloneDeMix has high specificity for detecting early mutations in tumor progression; these early mutations would be good candidates for disease driver genes and targeted therapies.

## Acknowledgments

This work was supported by the Ministry of Science and Technology [MOST 105-2118-M-007-001].

## Supporting Information

**S1 Table. Reference list of cell-migration-related genes**

**S2 Table. List of reference genes in 11q13.3**

**S1 Fig. Copy number estimation results**

The estimated copy number states for the exons across the genome are presented in different colors. Light blue and red represent the deletion and amplification events, respectively. Black indicates no copy number changes.

**S1 Software**.

Software S1 is an R package called “CloneDeMix” that implements subclonal copy number decomposition and it is available at https://github.com/AshTai/CloneDeMix.

